# Complete mapping of viral escape from neutralizing antibodies

**DOI:** 10.1101/086611

**Authors:** Michael B. Doud, Scott E. Hensley, Jesse D. Bloom

## Abstract

Identifying viral mutations that confer escape from antibodies is crucial for understanding the interplay between immunity and viral evolution. Here we quantify how every amino-acid mutation to influenza hemagglutinin affects neutralization by monoclonal antibodies targeting several antigenic regions. Our approach involves creating all replication-competent protein variants of the virus, selecting these variants with antibody, and using deep sequencing to identify enriched mutations. These high-throughput measurements are predictive of the effects of individual mutations in traditional neutralization assays. At many residues, only some of the possible mutations escape from an antibody. For instance, at a single residue targeted by two different antibodies, we identify some mutations that escape both antibodies and other mutations that escape only one or the other. Therefore, our approach maps how viruses can escape antibodies with mutation-level sensitivity, and shows that only some mutations at antigenic residues actually alter antigenicity.

## INTRODUCTION

Host immunity drives the evolution of many viruses. For instance, selection from antibodies leads to the fixation of amino-acid mutations in the hemagglutinin (HA) of human seasonal influenza A virus at a rate of over two substitutions per year (Smith et al., 2004; Bedford et al., 2015). However, it is difficult to predict *a priori* how any given viral mutation will contribute to immune escape.

A classic approach for identifying immune-escape mutations is to select individual viral mutants that are resistant to neutralization by antibodies. For instance, escape-mutant selections with a panel of monoclonal antibodies were used to broadly define several major antigenic regions of influenza HA (Webster and Laver, 1980; Gerhard et al., 1981; Caton et al., 1982). However, each such experiment typically only selects one of the potentially many mutations that can escape an antibody, with a strong bias towards whichever mutations happen to be prevalent in the initial viral stock. Therefore, escape-mutant selections provide an incomplete picture of the ways that a virus can escape an antibody.

A complete structural definition of the physical interface between an antibody and antigen can be obtained using methods such as X-ray crystallography. However, such structural definitions do not reveal which mutations actually ablate antibody recognition. For instance, alanine scanning has shown that mutations at only a fraction of the sites in the antibody-antigen interface actually disrupt antibody binding (Jin et al., 1992; Muller et al., 1998; Dall’Acqua et al., 1998; Pons et al., 1999), a “hot spot” phenomenon observed in protein-protein interfaces more generally (Cunningham and Wells, 1993; Clackson and Wells, 1995; Bogan and Thorn, 1998). Therefore, even crystal structures of antibodies bound to viral proteins do not delineate which mutations are functionally relevant for immune escape.

Here we use massively parallel experiments to map all amino-acid mutations that enable influenza virus to escape from several neutralizing antibodies. Our approach involves imposing antibody selection on virus libraries generated from all amino-acid mutants of HA, and then using deep sequencing to quantify how every mutation affects antibody neutralization of actual replication-competent influenza virus. Our comprehensive mapping of viral escape reveals remarkable mutation-level idiosyncrasy in antibody escape: for instance, at many residues only some of the possible amino-acid mutations confer escape, and two antibodies targeting the same residue elicit unique profiles of escape mutations. Mutational antigenic profiling therefore provides a high-resolution map of viral antibody escape mutations.

## RESULTS

### Mutational antigenic profiling reproducibly measures the effects of all HA mutations on antibody neutralization

To measure how all amino-acid mutations to HA affect virus escape from neutralizing antibodies, we developed the mutational antigenic profiling strategy shown in Figure 1. Viruses carrying all amino-acid mutations to HA compatible with viral replication are incubated with or without a neutralizing antibody, and then used to infect cells. Deep sequencing is then used to measure the frequency of each mutation among the viral variants that infected cells in either the presence or absence of antibody. We quantify the *differential selection* for each mutation as the logarithm of its enrichment in the antibody-treated virus library relative to the no-antibody control, and display these data as shown in Figure 1B. In the analysis that follows, we only consider mutations with positive differential selection.

**Figure 1:**
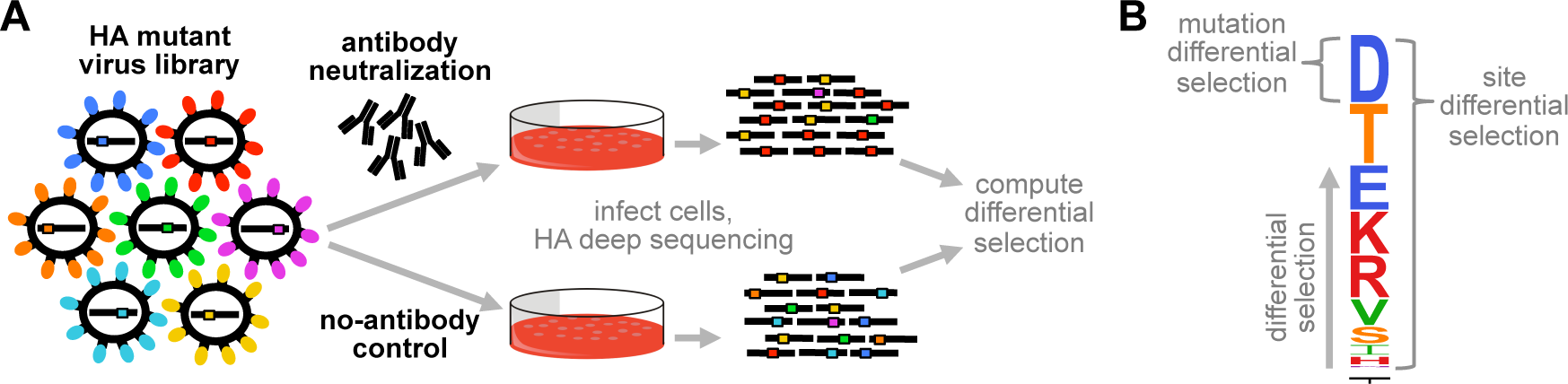
Overview of mutational antigenic profiling. **(A)** Libraries of influenza viruses carrying all mutants of HA that support viral replication are incubated with or without antibody, and used to infect cells. Viral RNA is extracted from cells and deep sequenced to quantify the frequency of each mutation in the antibody-selected and no-antibody control samples. **(B)** Differential selection is defined as the logarithm of the enrichment of each mutation in the antibody-selected sample versus the no-antibody control. In the logo plots, the height of each letter is proportional to the differential selection for that amino-acid. The site differential selection is the total height of the logo stack at that site (the sum of all mutation differential selection values). Only positive differential selection (corresponding to mutations enriched by antibody selection) is shown. In all logo plots, letters are colored by physicochemical properties of amino-acids as explained in the Methods.

We began by applying this approach to map all escape mutants in the A/WSN/1933 (H1N1) HA from a monoclonal antibody (H17-L19) targeting the Ca2 antigenic region (Gerhard et al., 1981). We have previously described virus libraries of all replication-competent amino-acid variants of A/WSN/1933 HA (Thyagarajan and Bloom, 2014; Doud and Bloom, 2016). We performed three full biological replicates of the mutational antigenic profiling using three independently generated mutant virus libraries, as well as a technical replicate of the profiling with one of the virus libraries (Figure 2A). The rationale for performing biological and technical replicates was to evaluate experimental noise arising both from variability in the initial virus libraries and from stochasticity in repeated selections on the same library.

**Figure 2:**
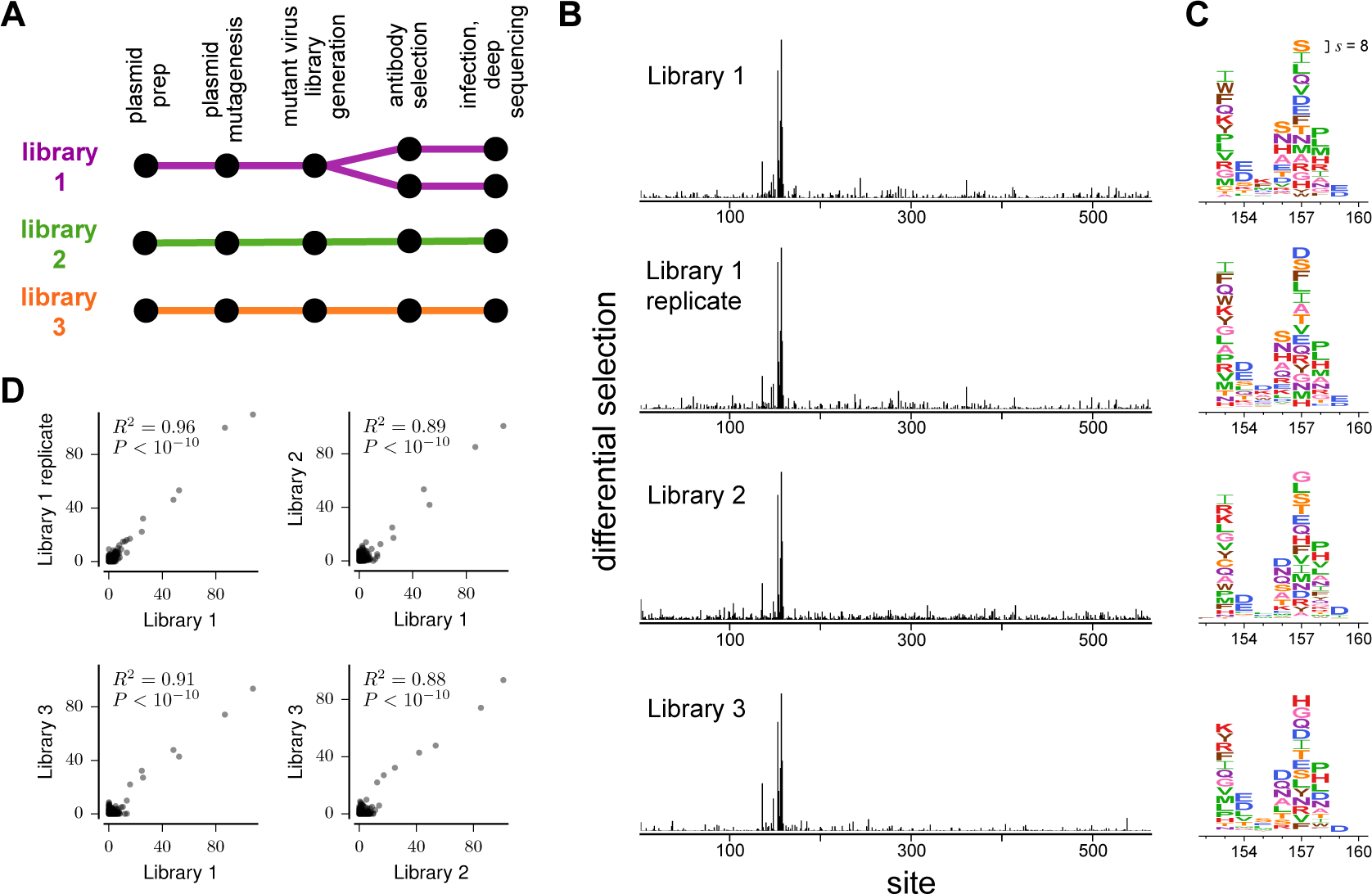
Mutational antigenic profiling with antibody H17-L19 is highly reproducible. **(A)** We performed three biological replicates and one technical replicate. **(B)** Site differential selection across HA is focused on the same subset of sites in all biological and technical replicates. **(C)** A zoomed-in view of the selection on the core of the antibody epitope. The height of each letter is proportional to the differential selection for that amino-acid. The same scale is used in all panels of (B) and (C). The scale bar in the upper-right of (C) shows the letter height for a mutation with differential selection of 8, corresponding to a 2^8^ = 256-fold enrichment by antibody selection. Residues are numbered sequentially beginning with the initiating methionine; conversions to other numbering schemes are in Supplementary file 1. **(D)** The site differential selection across all HA sites is highly correlated among replicates. Each point represents selection at one site in HA; correlation coefficients are Pearson’s R. Data is shown for selections with antibody H17-L19 at a concentration of 10 *μ*g/ml.

In each replicate experiment, the antibody exerted strong differential selection for mutations at a handful of sites, and little selection on the rest of HA (Figure 2B). Figure 2C shows the selection for individual amino-acid mutations in a short region in HA containing the majority of the epitope. Visual inspection of Figure 2B,C reveals consistent selection across technical and biological replicates. Statistical analysis confirms that the site differential selection is strongly correlated among all replicates (Figure 2D).

We next asked how the differential selection depended on the concentration of antibody used. Figure 2 shows the results of mutational antigenic profiling at an antibody concentration where the virus libraries retained only 0.3% of their total infectivity. We performed additional experiments using several dilutions of antibody that spanned a 20-fold range. Figure 3A shows the selection averaged across replicates at each antibody concentration. As expected, there is minimal differential selection when comparing replicate no-antibody controls. At progressively higher antibody concentrations, differential selection increases at most sites in the antibody epitope, while noise at other sites remains similar to that in the no-antibody control. However that the increase in differential selection with antibody concentration is not entirely uniform across sites (Figure 3A). Understanding the exact determinants of how a mutation’s differential selection depends on antibody concentration is an interesting area for future work. However, Figure 3B shows that despite these complexities, the sites of greatest differential selection are similar across antibody concentrations, indicating that the identification of escape mutations does not strongly depend on antibody concentration within the 20-fold range tested here.

**Figure 3:**
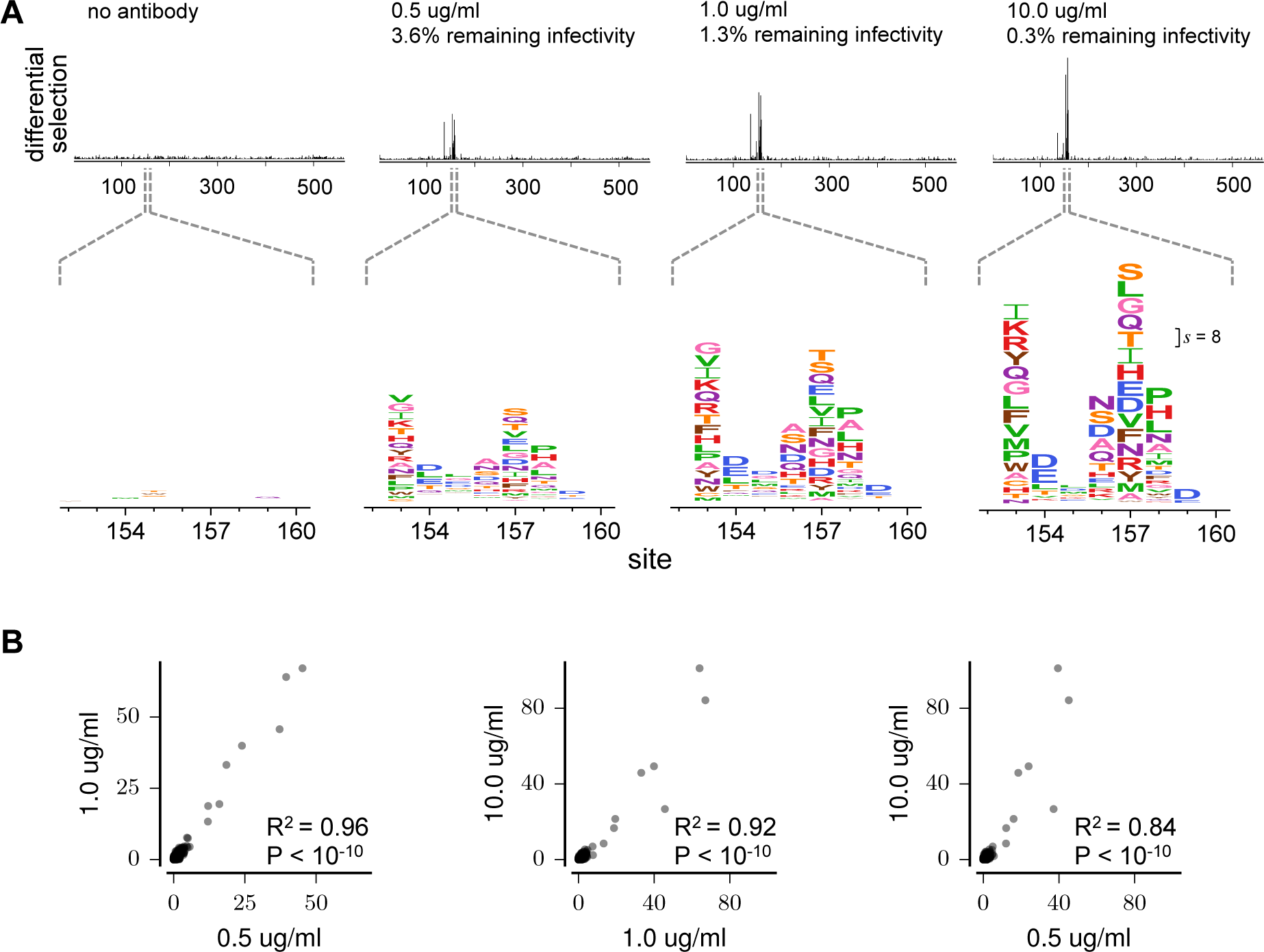
Differential selection by H17-L19 at different antibody concentrations. (A) Differential selection increases with progressively higher antibody concentrations. The top plots show site differential selection across the entire HA gene; the bottom plots show the core of the epitope. All horizontally-aligned plots use the same scale. The scale bar in the right-most logo plot shows the letter height for a mutation with differential selection of 8, corresponding to a 2^8^ = 256-fold enrichment by antibody selection. The “no antibody” differential selection is computed between two replicate experiments on a single library. (B) Site differential selection is correlated between antibody concentrations, although the strength of selection generally increases at most sites with higher antibody concentration. Each point represents selection at one site in HA; correlation coefficients are Pearson’s R. The data for each concentration is the average across the three mutant virus library replicates.

Overall, these results confirm that mutational antigenic profiling reproducibly identifies the HA mutations that confer escape from the monoclonal antibody H17-L19. The identified sites of selection are robust across replicate virus libraries and concentrations of antibody.

### Complete mapping of escape mutations for four antibodies

We next extended the mutational antigenic profiling to three more antibodies. We performed selections with each antibody at concentrations at which the virus libraries retained 0.1 to 0.2% of their infectivity (Supplementary table 1). Each antibody exerted strong selection at a small number of residues in HA. Figure 4A shows the site differential selection along the entire length of HA, while Figure 4B shows detailed mutation-level selection at key positions in the antibody epitopes. We again performed three full biological replicates with each antibody, and the results were again highly reproducible among replicates (Supplementary figure 1).

**Figure 4:**
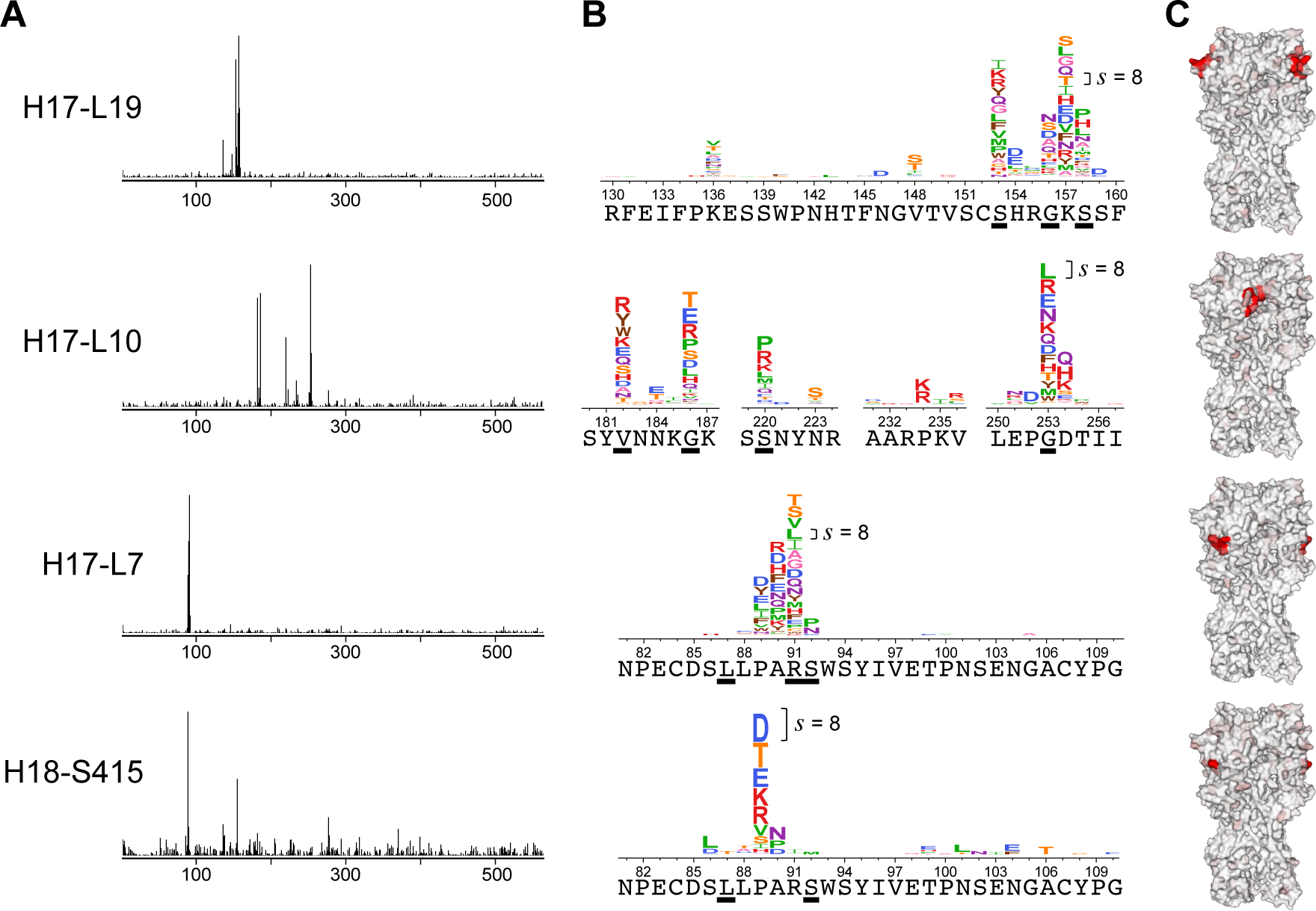
Mutational antigenic profiling of four different antibodies. **(A)** Each antibody exerts a different profile of selection on HA. **(B)** Zoomed in view of the most strongly selected sites for each antibody. The wild-type amino acid at each site is shown under the logoplots. Sites where escape mutations were selected by Gerhard et al. (1981) and Caton et al. (1982) are underlined. **(C)** The functional epitope for each antibody is visualized on HA’s three-dimensional structure (PDB 1RVX, Gamblin et al., 2004). Each site is colored from white to red based on the differential selection for the most strongly selected mutation at that site for each antibody. Red indicates strong differential selection. All structures show trimeric HA in the same orientation (the epitope is visible for two of the three monomers for H17-L19, H17-L7, and H18-S415). The y-axis scale is set separately for each antibody. The scale bar in each logo plot shows the letter height for a mutation with differential selection of 8, corresponding to a 2^8^ = 256-fold enrichment by antibody selection. The data for each antibody is the average across the three mutant virus library replicates. Supplementary table 1 shows the percentage of each mutant virus library remaining infectious after antibody neutralization in each replicate experiment. Site differential selection is highly correlated among the replicates for each antibody (Supplementary figure 1). A zoomed in structural view of the most strongly selected regions of HA for each antibody is shown in Supplementary figure 2. Supplementary table 2 lists residues where escape mutations were previously identified by classic escape-mutant selections. Logo plots and files providing differential selection values across the entire gene are in Supplementary file 3 and Supplementary file 4.

For each antibody, the sites of strongest differential selection were clustered in a single surface-exposed patch on HA’s structure that is presumably within the antibody-binding footprint (Figure 4C and Supplementary figure 2). The four antibodies target three antigenic regions: H17-L19 targets Ca2, H17-L10 targets Ca1, and H17-L7 and H18-S415 both target Cb (Gerhard et al., 1981; Caton et al., 1982). As expected, H17-L19 and H17-L10 exert strong selection on entirely distinct sets of residues, but H17-L7 and H18-S415 exert selection on similar sets of residues in the Cb antigenic region. For three of the antibodies, the strongly selected residues are within short contiguous stretches of primary amino-acid sequence, but for H17-L10 the strongly selected residues are distributed across 70 residues of HA’s primary sequence.

### Comparison to classical escape-mutant selections

The antigenicity of H1 HA was originally characterized in a classic set of experiments that selected individual viral escape mutants with a panel of mouse monoclonal antibodies (Gerhard et al., 1981; Caton et al., 1982). These experiments identified a handful of mutations that ablated binding by each antibody. All four antibodies used in our study are from the original panel used by Gerhard et al. (1981). We expected that the sites of differential selection identified in our complete mapping of viral escape mutants would include the previously identified mutations.

Indeed, there is strong overlap between the sites identified by our mutational antigenic profiling and the mutations selected in the classic escape-mutant selections (see underlined residues in Figure 4B). However, we also identified numerous additional escape mutations at those and other sites. In some cases, the sites that we found to be under the strongest differential selection were not identified at all in the classic escape-mutant selections. For instance, as shown in Figure 4, the classic escape-mutant selections failed to identify site 157 for H17-L19, site 89 for H17-L7, and site 89 for H18-S415 (throughout this manuscript, residues are numbered sequentially beginning at the N-terminal methionine; conversions to other numbering schemes are in Supplementary file 1). Presumably mutations at these sites were not uncovered in the escape-mutant selections because each such selection only finds one mutation, with a strong bias towards those that arise from single-nucleotide changes that are prevalent in the viral stock. In contrast, our approach simultaneously examines all possible amino-acid mutations. Therefore, in a single set of experiments, we have completed the mapping of escape mutants to four antibodies begun 35 years ago by Gerhard et al.(1981).

### Validating the mutation-level sensitivity of mutational antigenic profilling

The results described above were obtained from high-throughput experiments that used deep sequencing to examine tens of thousands of viral variants in parallel. Are these results predictive of the antigenic effects of mutations examined individually? To answer this question, we tested some of our key findings with neutralization assays on individual viral mutants. To do this, we used site-directed mutagenesis to introduce single amino-acid mutations into the HA gene, generated viruses by reverse genetics, and performed GFP-based neutralization assays.

One striking observation from our mutational antigenic profiling is that at some residues, only a few of the possible amino-acid mutations are strongly selected by any given antibody. For instance, at HA residue 154, the H17-L19 antibody exerts strong selection only for mutations H154E and H154D, both of which introduce a negatively charged amino acid (Figure 4B, Figure 5A). To validate this result, we generated viruses carrying the H154E mutation or a mutation to alanine (H154A), which our mutational antigenic profiling did not find to be under differential selection. Neutralization assays confirmed that the H154E mutant completely escaped at all antibody concentrations tested, while the H154A mutant was as sensitive to antibody as wild-

**Figure 5:**
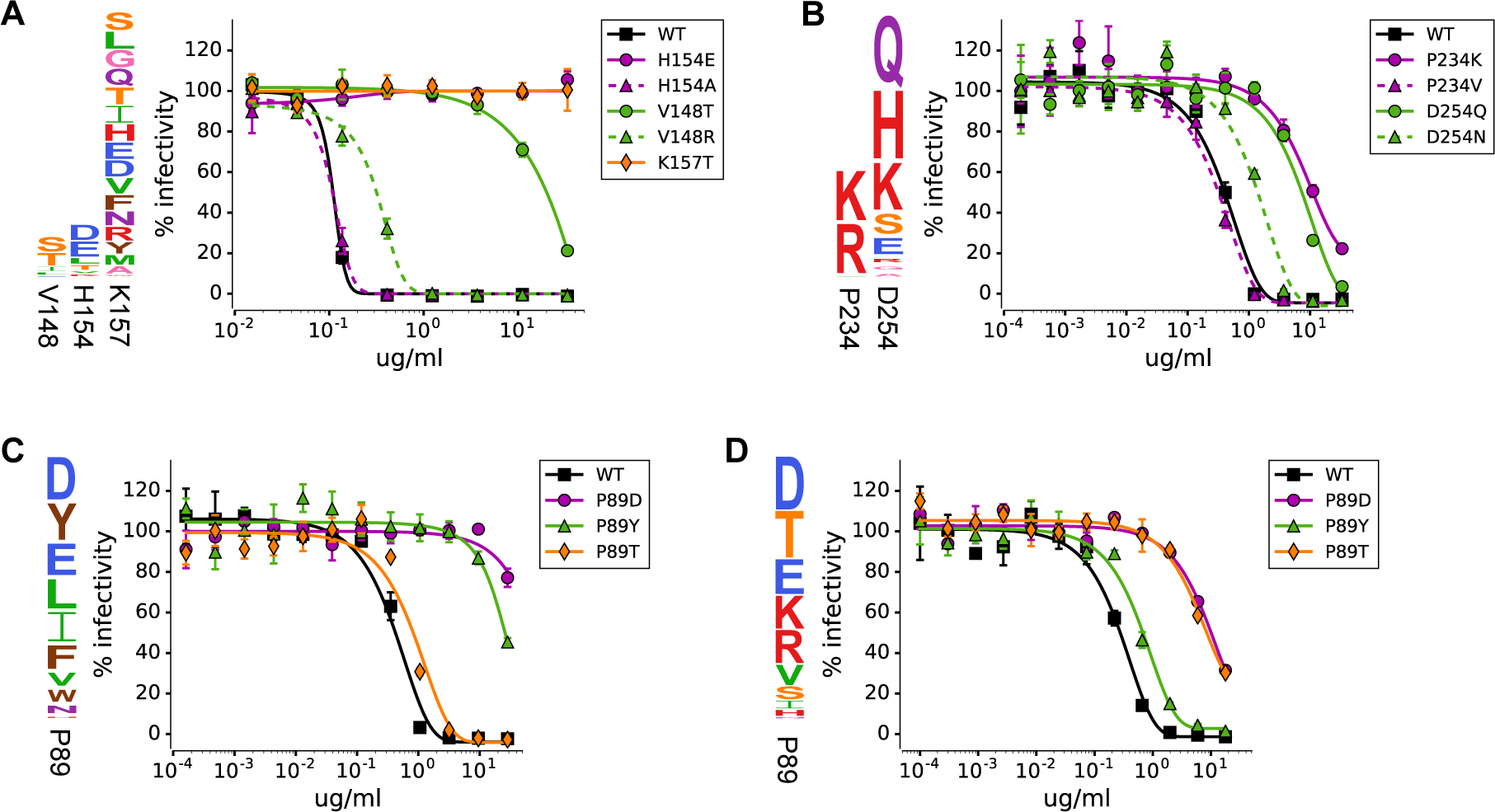
Validation of the mutational antigenic profiling with neutralization assays on viruses carrying single mutations. In each panel, the logo plot at left shows the results of the mutational antigenic profiling at sites selected for validation, and the graph at right shows the results of the neutralization assays. Note how in many cases, only some of the amino-acid mutations at a site escape from any given antibody. The antibodies shown in each panel are: **(A)** H17-L19, **(B)** H17-L10, **(C)** H17-L7, **(D)** H18-S415.type (Figure 5A). Therefore, a more limited method such as alanine scanning would not have identified residue 154 as a site of escape mutations. This finding demonstrates the importance of assaying all amino-acid mutations if the goal is to comprehensively map sites of escape.

Another example of mutation-level sensitivity is HA residue 148, where antibody H17-L19 only selects for mutations to serine and threonine (Figure 4B). Both of these mutations (V148T and V148S) introduce a motif (N-X-S/T) that potentially leads to glycosylation of the asparagine at site 146. To validate that only some mutations at site 148 enable escape, we generated the V148T mutant as well as another mutant (V148R) that does not introduce a glycosylation motif. As expected, V148T dramatically reduced the virus’s sensitivity to the antibody, whereas V148R only had a small effect (Figure 5A).

Our high-throughput results suggested similar mutation-level sensitivity in escape from antibody H17-L10. At residue 234, the mutational antigenic profiling identified strong differential selection only for mutations to the positively charged amino-acid residues lysine and arginine (Figure 4B). We generated a virus carrying one of these mutations (P234K) as well as a virus carrying another mutation at the same residue (P234V) that our results predicted would have no effect. Neutralization assays confirmed that the P234K mutation escaped H17-L10, while the P234V mutation caused no change in antibody sensitivity (Figure 5B). Interestingly, in HA’s structure, site 234 is on a neighboring protomer relative to all the other mutations strongly selected by H17-L10 (Supplementary figure 2). Our finding that escape mutations from H17-L10 cross the HA trimer interface is consistent with the fact that this antibody only recognizes trimeric HA (Magadán et al., 2013). Escape mutations at such epitopes are discernible because mutational antigenic profiling uses actual viruses that display intact HA; such conformational epitopes might not be properly displayed in the modified forms of viral glycoproteins often used in other high-throughput methods such as phage and yeast display.

Overall, these results indicate the power of mutational antigenic profiling to map residues where only a few specific amino-acid mutations lead to escape from antibody. Because this approach examines HA in its native context on influenza virions, it can comprehensively map escape mutations even in complex conformational epitopes.

### Unique repertoires of escape mutations from two antibodies targeting the same site in HA

Two of the antibodies used in our study (H17-L7 and H18-S415) target the same antigenic region of HA, with residue 89 under strong selection from both antibodies (Figure 5C,D). Do these antibodies select the same or different amino-acid mutations at this residue? Our mutational antigenic profiling suggests that both antibodies select mutations to negatively charged amino acids (P89D and P89E; Figure 5C,D). However, each antibody also selects a unique set of additional mutations, such as P89Y for H17-L7 and P89T for H18-S415.

We generated viruses containing the P89D, P89Y, or P89T mutations and tested their sensitivity to both antibodies using neutralization assays. In agreement with the mutational antigenic profiling, the P89D mutant escaped both antibodies, but P89Y only escaped from H17-L7 and P89T only escaped from H18-S415 (Figure 5C,D). Thus, when two antibodies target the same site, there can be both common and antibody-specific routes of escape. Characterizing antibody escape at the level of protein sites therefore only provides a partial picture of antigenicity. A complete understanding of escape requires consideration of every mutation at every site.

## DISCUSSION

We have used a new high-throughput approach to completely map the amino-acid mutations that enable influenza virus to escape from several neutralizing antibodies. Our approach is conceptually similar to recent methods that couple deep sequencing with phage or yeast display assays for antibody binding (Kowalsky et al., 2015; Adams et al., 2016; Gaiotto and Hufton, 2016). But whereas those methods are limited to selecting for binding to antigens expressed in bacteria or yeast, our approach selects for actual neutralization in the context of replication-competent virus. Our experiments therefore measure the phenotype truly relevant to virus evolution: whether a mutation enables a virus to escape neutralization by an antibody.

Our approach also bears similarities to the classic method of selecting individual viral escape mutants. However, classic escape-mutant selections are low throughput and rely on the occurrence of *de novo* mutations in a viral stock. Therefore, like evolution itself, such selections are stochastic, and only identify one of potentially many escape mutations. In contrast, our massively parallel experiments simultaneously examine all amino-acid mutations, thereby eliminating stochasticity and allowing us to completely define the possible antibody-escape mutations.

The most striking finding from our work is the exquisite mutation level-sensitivity of antibody escape. For each of the four antibodies, our high-throughput experiments identified residues in HA where only some of the possible amino-acid mutations conferred escape. For each antibody, we used neutralization assays to validate specific residues where one mutation mediated escape whereas another had little or no effect. In some cases, this mutation-level sensitivity is easy to rationalize: we found examples where escape required mutations that introduce glycosylation motifs or change the charge of the amino-acid sidechain. But in other cases, the effects are not only difficult to rationalize but depend on the specific antibody. For instance, we identified a residue targeted by two antibodies where a mutation that escaped the first antibody had no effect on the second and vice versa. Previous studies have distinguished between an antibody’s “functional epitope” and physical footprint based on the observation that binding is disrupted by mutations at only some residues that contact the antibody (Jin et al., 1992; Muller et al., 1998; Dall’Acqua et al., 1998; Pons et al., 1999). Our findings extend this concept by showing that even within the functional epitope, only certain mutations actually mediate escape.

These results underscore the shortcomings of thinking about viral antigenic evolution purely in terms of antigenic sites. For instance, many approaches to forecast and model influenza virus evolution are based on partitioning HA into antigenic and non-antigenic sites (Bush et al., 1999; Łuksza and L¨assig, 2014; Neher and Bedford, 2015). However, our work shows that for any individual antibody, it is important to consider the exact amino-acid mutation as well as the site at which it occurs. Future application of our massively parallel mutational antigenic profiling approach to contemporary viral strains and antibodies could aid in prospectively identifying the antigenic consequences of viral mutations on a vastly more comprehensive scale than has previously been possible (Li et al., 2016).

## Materials and Methods

### Mutant virus libraries

The influenza virus mutant libraries used in this study are those described in Doud and Bloom (2016). As detailed in that reference, reverse-genetics plasmids (Neumann et al., 1999) encoding HA gene were mutagenized at the codon level using the protocol described in Bloom (2014). These plasmid codon-mutant libraries were then used to generate libraries of replication-competent influenza viruses using a helper-virus approach that reduced the bottlenecks associated with standard reverse genetics. The virus libraries were then passaged at low MOI to create a genotype-phenotype link between the HA protein on a virion’s surface and the gene that it carries. The viral titers in these libraries were determined by TCID_50_ (50% tissue culture infectious dose) in MDCK-SIAT1 cells. Three fully independent virus libraries were generated beginning with independent plasmid mutant libraries as outlined in Figure 2A. It was these low-MOI passaged virus libraries, which are characterized in detail in Doud and Bloom (2016), that formed the starting point for the antibody selections described in the current work.

### Antibodies

The antibodies used in this study were originally isolated from mice as described in Gerhard et al. (1981) and Caton et al. (1982). Note that in these older papers, two different naming schemes are used for the same antibodies: H17-L19 was also called Ca3; H17-L10 was also called Ca6; H17-L7 was also called Cb15; H18-S415 was also called Cb5. Antibodies secreted by H17-L19, H17-L10, H17-L7, and H18-S415 hybridoma cell lines were purified using PureProteome A/G coated magnetic beads (Millipore).

### Mutant virus selections with antibody

For the selections outlined in Figure 1, we began by diluting each virus library in influenza growth media (Opti-MEM supplemented with 0.3% BSA, 100 U of penicillin/ml, 100 *μ*g of streptomycin/ml, and 100 *μ*g of calcium chloride/ml) to a concentration of 1 × 10^6^ TCID_50_ per ml. Monoclonal antibody was also diluted in influenza growth media to a concentration twice that intended for use the selection. The virus library was then neutralized by mixing 1 ml of diluted virus with 1 ml of diluted antibody to give the final antibody concentrations listed in Supplementary table 1. This virus-antibody mixture was then incubated at 37°C for 1.5 hours. No-antibody controls were “mock-neutralized” in parallel by substituting influenza growth media for the diluted antibody. At the same time, serial ten-fold dilutions of mutant virus library were made from the 1 × 10^6^ TCID_50_ per ml virus stock to be used as a standard curve to measure infectivity. These dilutions represented 10%, 1%, 0.1%, 0.01%, and 0.001% of the 1 × 10^6^ TCID_50_ dose of library used in neutralizations.

The viral samples were then added to cells to allow infection by non-neutralized virions. We used MDCK-SIAT1 cells that had been plated four hours prior to infection in D10 media (DMEM supplemented with 10% heat-inactivated FBS, 2 mM L-glutamine, 100 U of penicillin/ml, and 100 μg of streptomycin/ml) at 2.5 × 10^5^ cells per well in 6-well dishes. For the infections, we aspirated off the existing D10 media and added the 2 ml of viral sample. Duplicate infections were used for each point on the standard curve of serially diluted virus. After two hours, media in each well was then changed to 2 ml WSN growth media (Opti-MEM supplemented with 0.5% heat-inactivated FBS, 0.3% BSA, 100 U of penicillin/ml, 100 μg of streptomycin/ml, and 100 μg of calcium chloride/ml) after rinsing cells once with PBS to remove residual virus in the supernatant.

Twelve hours later, RNA was isolated from the cells in each well using a Qiagen RNEasy Plus Mini kit by aspirating media, adding 350 μl buffer RLT freshly supplemented with β-mercaptoethanol, slowly pipetting several times to lyse cells, transferring the lysate to a RNase-free microfuge tube, vortexing for 20 seconds to homogenize, and proceeding with the manufacturer’s suggested protocol, eluting in 35 μl of RNase-free water.

We estimated the percent remaining infectivity in the neutralized samples using qRT-PCR and a standard curve created using the infections with 10-fold serial dilutions of the virus libraries to give the estimates in Supplementary table 1. For the qPCR, primers WSN-NP-qPCR-F (5’-GCAACGGCTGG TCTGACTCACA-3’) and WSN-NP-qPCR-R (5’-TCCATTCCTGTGCGAACAAG-3’) were used to amplify influenza nucleoprotein (NP) to quantify viral infectivity, and primers 5’-canineGAPDH (5’-A AGAAGGTGGTGAAGCAGGC-3’) and 3’-canineGAPDH (5’-TCCACCACCCTGTTGCTGTA-3’) were used to quantify canine GAPDH to correct for small differences in total RNA amounts. qRT-PCR was performed using Applied Biosystems PowerSYBR green RNA-to-Ct 1-step kit, with 40 ng of RNA in each 20 μl reaction, cycling conditions of 48°C for 30 minutes, 95°C for 10 minutes, and 40 cycles of: 95°C for 15 sec, 58°C for 1 min with data acquisition. All samples were measured in duplicate, and each assay included no-reverse-transcriptase controls. Linear regression of the relationship between the log(infectious dose) and the mean difference in Ct between NP and GAPDH was used to interpolate the remaining infectious dose of each antibody-neutralized sample, expressed as a percentage of the 1 × 10^6^ TCID_50_ used in each neutralization.

### Deep sequencing and quantification of mutation frequencies

To prepare deep sequencing libraries, HA genes were amplified from the RNA isolated from infected cells by reverse transcription with AccuScript Reverse Transcriptase (Agilent 200820) using HA-specific primers WSN-HA-for (5’-AGCAAAAGCAGGGGAAAATAAAAACAAC-3’) and WSN-HA-rev (5’-AGTAGAAACAAGGGTGTTTTTCCTTATATTTCTG-3’). PCR amplification of HA cDNA and Illumina sequencing library preparation was then carried out using the barcoded subamplicon sequencing protocol in Doud and Bloom (2016), which was in turn inspired by the approach in Wu et al. (2014). The only change made to the protocol in Doud and Bloom (2016) was that in order to more effectively spread sequencing depth across samples based on the expected diversity of mutations in each sample, the number of uniquely-barcoded single stranded variants used as template for round 2 PCR was 5 × 10^5^ to 7 × 10^5^ for the no-antibody control samples, and 1.5 × 10^5^ for the antibody-neutralized samples. Sequencing libraries with unique indices for each experimental sample were pooled and sequenced on an Illumina HiSeq2500 using 2 x 250 bp paired-end reads in rapid-run mode.

The frequency of each mutation in each sample was determined by using dms tools (Bloom, 2015, http://jbloomlab.github.io/dms_tools/), version 1.1.20, to align subamplicon reads to a reference HA sequence, group barcodes to build consensus sequences, and quantify mutation counts at every site in the gene for each experimental sample.

### Computation of differential selection

We computed the extent that each mutation is enriched by each antibody selection by comparing mutation counts in each antibody-treated sample to mutation counts from the matching no-antibody control sample. We compute the *differential selection* on each mutation between the two samples (Figure 1B) using the analysis framework introduced in Ashenberg et al. (2016), with slight modifications to account for PCR and sequencing errors. The error rate *ε*_r,x_ at each site *r* for codon *x* is estimated from the apparent frequency of that mutation in our previously described sequencing of HA from wild-type plasmid using barcoded-subamplicon Illumina sequencing (Doud and Bloom, 2016). Specifically, the error rate was calculated as:

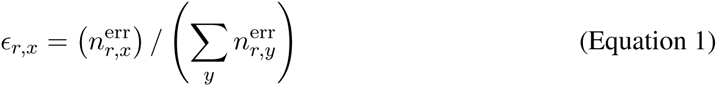

where 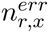 is the number of counts of codon *x* at site *r* in the wild-type plasmid sequencing library. We then adjusted the observed counts 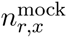 and 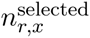 for codon *x* at site *r* in the mock selected and antibody selected samples, respectively, to the error-corrected counts 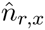 for each sample:

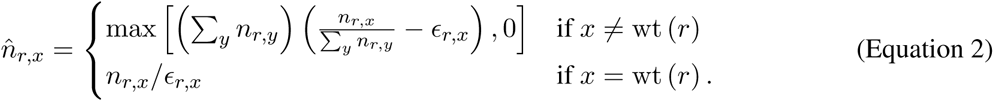

To convert from codon counts to amino-acid counts, we summed the error-adjusted counts for all codons encoding each amino acid *a* at site *r* to give the error-adjusted amino-acid counts 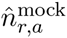 and 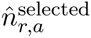 for the mock selected and antibody selected samples, respectively. We then computed the relative enrichment *E_r,a_* of amino acid *a* at site *r* as

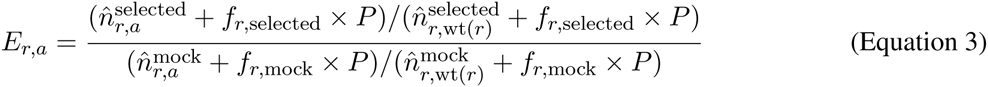

where wt (*r*) denotes the wildtype amino acid at site *r*, *P* is a pseudocount (set to 10 in our analyses), and *f_r,_*selected and *f_r,_*_mock_ give the relative depths of the selected and mock samples at site *r*:

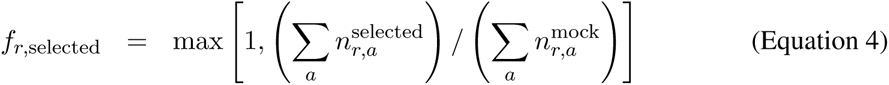

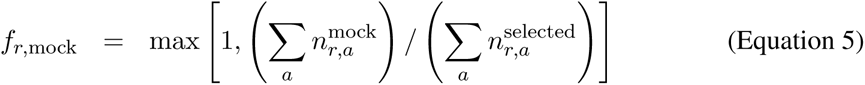

The reason for scaling the pseudocount by the library depth is that in the absence of such scaling, if the selected and mock samples are sequenced at different depths, the estimates of *E_r,a_* will tend to be systematically different from one even if the relative counts are the same in both conditions.

The mutation differential selection values are the logarithm of the enrichment values:

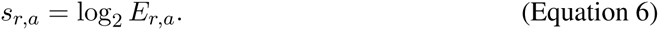

Mutations that confer escape from an antibody will have a larger relative frequency in the antibody-selected sample than the no-antibody control sample, and will thus have a large, positive differential selection. Therefore, we limited analysis to positive differential selection to identify antibody escape mutations. To summarize the differential selection at each site, we sum the mutation differential selection values *s_r,a_* over all amino-acids *a* with positive mutation differential selection and term this the positive site differential selection *s_r_* for site *r*:

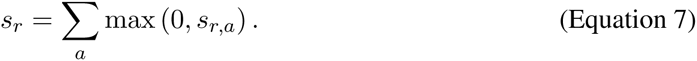

Logoplots visualizing differential selection display each amino acid with a height proportional to the mutation differential selection *s_r,a_*. Amino acid letter codes are colored based on the physiochemical properties of the amino-acid side chain: hydrophobic (V, L, I, M, P) are green, nucleophilic (S, T, C) are orange, small (A, G) are pink, aromatic (F, Y, W) are brown, amide (N, Q) are purple, positively-charged (H, K, R) are red, and negatively-charged (D, E) are blue.

The computer code to perform these differential selection analyses is incorporated in the dms_tools (http://jbloomlab.github.io/dms_tools/) software as the program `dms_diffselection.` The logoplots created by `dms_tools` are rendered with `WebLogo` (Crooks et al., 2004).

### GFP-based neutralization assays

We performed neutralization assays using viruses carrying GFP in the PB1 segment using the protocol described in Hooper and Bloom (2013). These GFP reporter viruses were generated using seven bidirectional reverse genetics plasmids (Hoffmann et al., 2000) encoding the PB2, PA, HA, NP, NA, M, and NS segments of A/WSN/1933 (kindly provided by Robert Webster of St. Jude Children’s Research Hospital), and a unidirectional reverse genetics plasmid pHH-PB1flank-GFP in which the coding sequence of PB1 is replaced by GFP (Bloom et al., 2010). Since these viruses carry GFP instead of PB1, they are grown in complementing 293T-CMV-PB1 and MDCK-SIAT1-CMV-PB1 cells that constitutively express the WSN PB1 protein.

For each HA mutation tested in the neutralization assay, the indicated amino-acid mutation was introduced into the WSN HA bidirectional reverse genetics plasmid by site-directed mutagenesis, and the HA sequence was verified by Sanger sequencing. To generate each mutant GFP-carrying virus, we transfected a co-culture of 293T-CMV-PB1 and MDCK-SIAT1-CMV-PB1 cells with the eight reverse genetics plasmids described above. For each transfection, 4 × 10^5^ 293T-CMV-PB1 and 4 × 10^4^ MDCK-SIAT1-CMV-PB1 per well were plated in 6-well plates in D10 media four hours prior to transfection. Each well received a transfection mixture of 100 μl DMEM, 3 μl BioT transfection reagent, and 250 ng of each of the eight reverse genetics plasmids. At 20 hours post-transfection, the media was changed to WSN neutralization media, which has low autofluorescence in the GFP channel (Medium 199 supplemented with 0.3% BSA, 100 U of penicillin/ml, 100 g of streptomycin/ml, 100 g of calcium chloride/ml, 25 mM HEPES, 0.5% FBS). At 72 hours post-transfection, culture supernatants were clarified by centrifugation at 2,000×g, aliquoted, and frozen at −80°C.

The GFP-carrying viruses were titered by flow cytometry in MDCK-SIAT1-CMV-PB1 cells. For this titering, cells were plated in 12-well plates at 1 × 10^5^ cells per well in WSN neutralization media. Four hours after plating, cells were infected with dilutions of viral supernatant. At 16 hours after infection, wells with approximately 1% of cells GFP-positive were analyzed by flow cytometry, and the fraction of GFP-positive cells was used to calculate the titer of infectious particles in each viral supernatant.

For the neutralization assays, monoclonal antibody was diluted down columns of a 96-well plate in WSN neutralization media. Three replicate dilution columns were used for each virus-antibody combination. Columns without antibody were used to measure maximal fluorescence in the absence of neutralization, and columns without cells were used to measure background fluorescence in viral supernatants, which we found to contribute more background fluorescence than cells alone. The GFP reporter viruses were diluted in WSN neutralization media to 1 × 10^3^ infectious particles per μl and 40 μl (4 × 10^4^ infectious particles) was added to each well. Plates were incubated at 37°C for 1.5 hours before adding 4 × 10^4^ MDCK-SIAT1-CMV-PB1 cells to each well. After 16 hours incubation at 37°C, GFP fluorescence intensity was measured on a Tecan plate reader using an excitation wavelength of 485 nm and an emission wavelength of 515 nm (12-nm slit widths). Percent of maximal infectivity was calculated by subtracting background fluorescence signal from all wells and dividing the signal from antibody-containing wells by the signal from corresponding wells without antibody.

### Data availability and source code

Deep sequencing data has been deposited at the Sequence Read Archive under biosample accession SAMN05789126. Supplementary file 2 contains an iPython notebook (and a static HTML version of this notebook) that performs all analyses of deep sequencing data. Logo plots of the differential selection by each antibody spanning the entire gene, averaged across the replicate libraries, are provided in Supplementary file 3. Files providing the mutation differential selection values, averaged across the replicate libraries for each antibody at each concentration tested, are in Supplementary file 4.

## Acknowledgments

We than Susanne L. Linderman for assistance with preparing the antibodies used in this study. We thank Bargavi Thyagarajan for assistance in developing initial ideas related to this project. This work was supported by grants R01GM102198 and R01AI127893 from the NIGMS and NIAID of the NIH. The research of JDB was supported in part by a Faculty Scholar grant from the Howard Hughes Medical Institute and the Simons Foundation. MBD was supported in part by training grant T32AI083203 from the NIAID of the NIH.

## Supplementary Material

**Supplementary figure 1:**
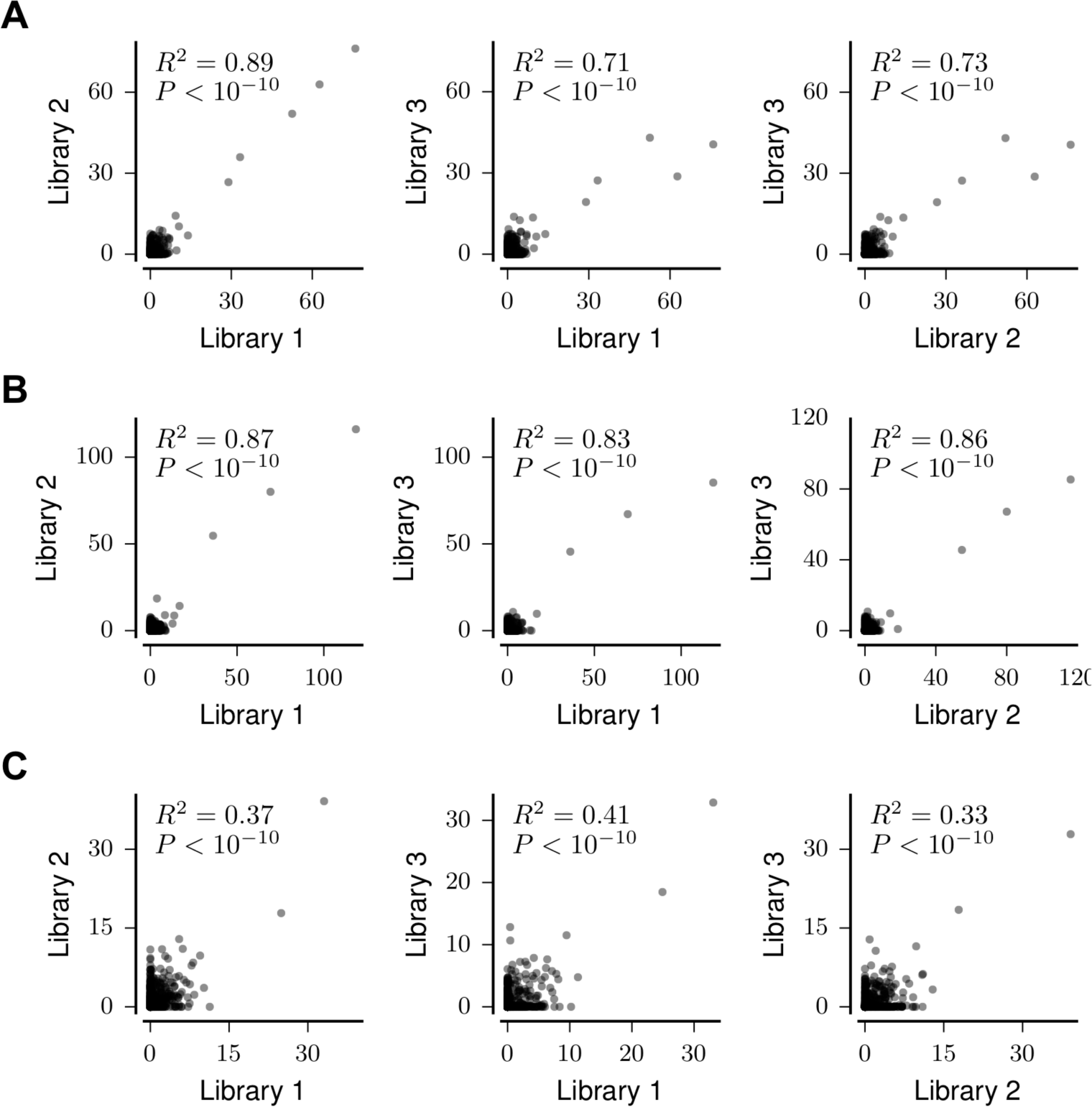
Positive site differential selection is highly correlated between full biological replicate measurements on independently generated mutant virus libraries for monoclonal antibodies **(A)** H17-L10, **(B)** H17-L7, and **(C)** H18-S415. Correlation coefficients are Pearson’s R.

**Supplementary figure 2:**
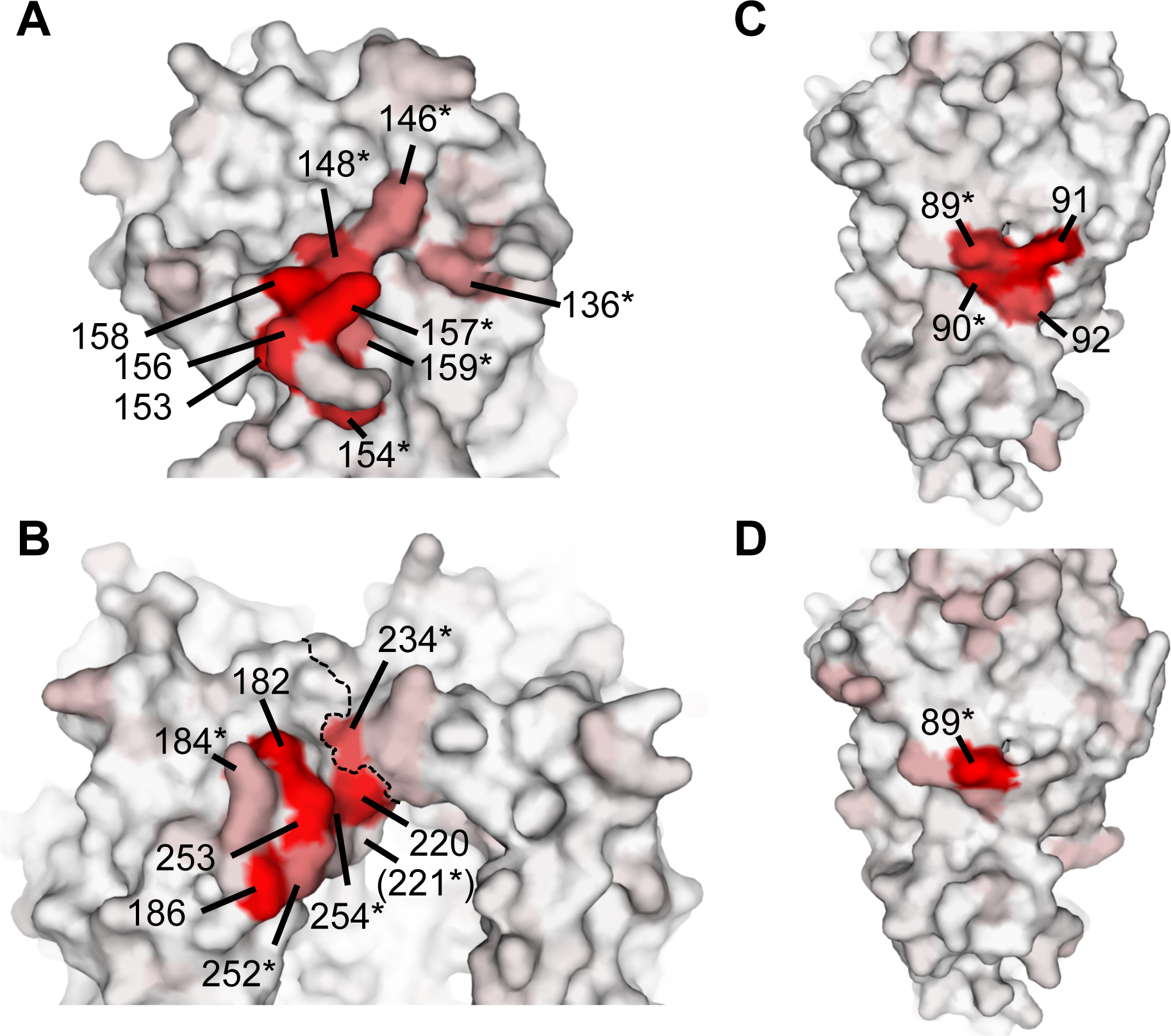
Detailed view of differential selection by each antibody projected onto HA’s structure. Each panel zooms into the relevant region of the structure shown in Figure 4C for that antibody. Residues are colored from white to red based on the differential selection for the most strongly selected mutation at that site for each antibody. Asterisks mark sites of strong differential selection which were not found in the original antigenic mapping of HA with that antibody (Gerhard et al., 1981; Caton et al., 1982). **(A)** H17-L19. **(B)** H17-L10. Strong differential selection at site 223 (not visible) results in putative glycosylation at site 221. The dashed line marks the boundary between two adjacent HA protomers. **(C)** H17-L7. **(D)** H18-S415.

**Supplementary table 1:** Percentage of each mutant virus library remaining infectious after antibody neutralization in each replicate selection experiment. Percent infectivity was measured by qRT-PCR of the influenza nucleoprotein gene and interpolated from a standard curve of infection prepared with serial dilutions of each virus library.

|  | library 1 | library 1 replicate | library 2 | library 3 |
| --- | --- | --- | --- | --- |
| H17-L19 0.5 $\mu\text{g/ml}$ | 2.1% | 2.5% | 4.9% | 5.4% |
| H17-L19 1 $\mu\text{g/ml}$ | 0.7% | 0.7% | 1.9% | 1.8% |
| H17-L19 10 $\mu\text{g/ml}$ | 0.3% | 0.2% | 0.4% | 0.4% |
| H17-L10 3 $\mu\text{g/ml}$ | 0.2% | | 0.2% | 0.2% |
| H17-L7 15 $\mu\text{g/ml}$ | 0.2% | | 0.1% | 0.1% |
| H18-S415 3 $\mu\text{g/ml}$ | 0.2% | | 0.1% | 0.2% |

**Supplementary table 2:**
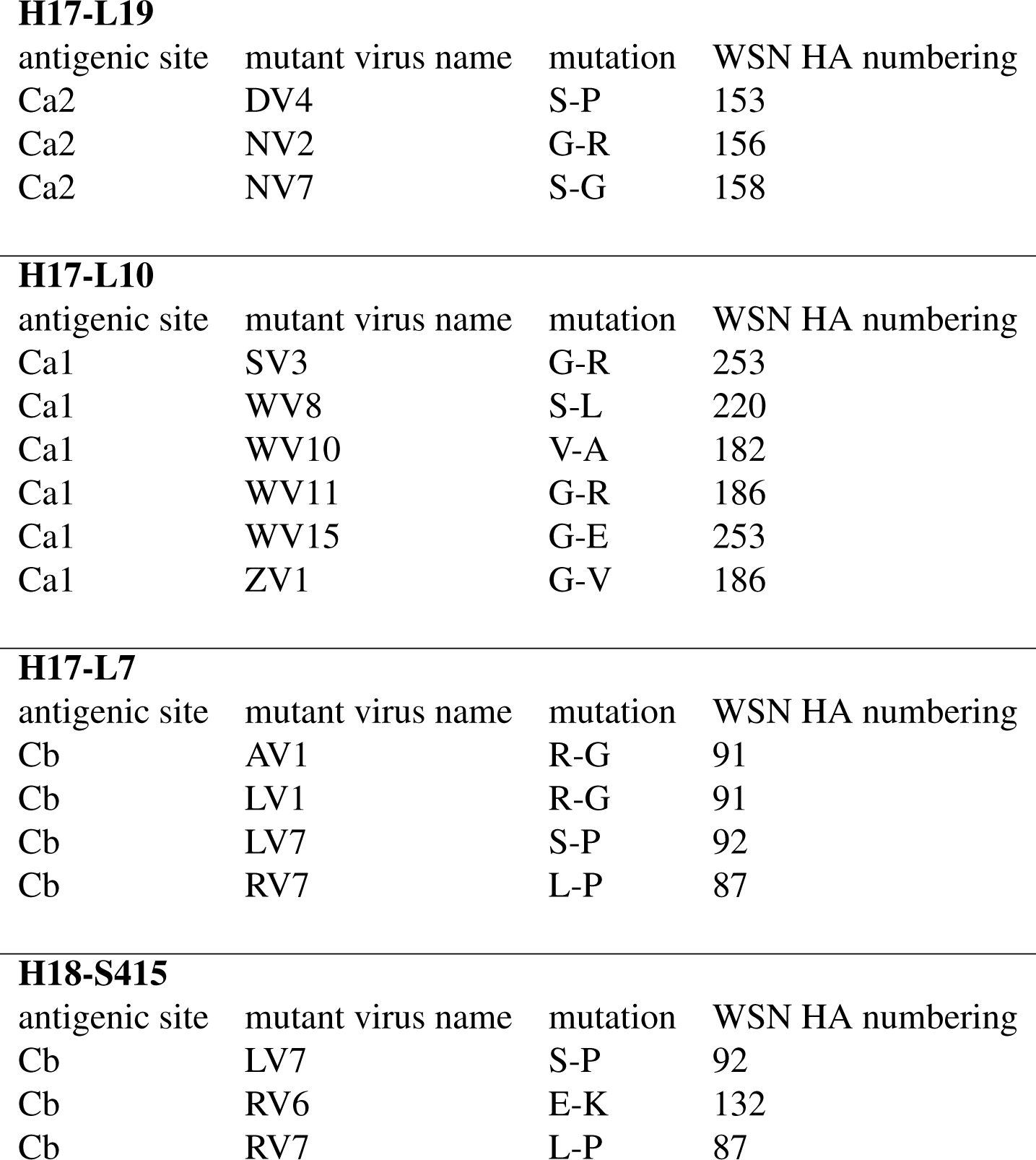
All mutations identified in the classic escape mutant selections by Gerhard et al. (1981) and Caton et al. (1982) with the four antibodies used in our study. Note that the classic experiments used the A/Puerto Rico/8/1934 (H1N1) virus, whereas our study used the A/WSN/1933 (H1N1) virus. In the older papers, multiple names were used to refer to the same antibody: H17-L19 was also called Ca3; H17-L10 was also called Ca6; H17-L7 was also called Cb15; H18-S415 was also called Cb5.

**H17-L19**
| antigenic site | mutant virus name | mutation | WSN HA numbering |
| --- | --- | --- | --- |
| Ca2 | DV4 | S-P | 153 |
| Ca2 | NV2 | G-R | 156 |
| Ca2 | NV7 | S-G | 158 |

**H17-L10**
| antigenic site | mutant virus name | mutation | WSN HA numbering |
| --- | --- | --- | --- |
| Ca1 | SV3 | G-R | 253 |
| Ca1 | WV8 | S-L | 220 |
| Ca1 | WV10 | V-A | 182 |
| Ca1 | WV11 | G-R | 186 |
| Ca1 | WV15 | G-E | 253 |
| Ca1 | ZV1 | G-V | 186 |

**H17-L7**
| antigenic site | mutant virus name | mutation | WSN HA numbering |
| --- | --- | --- | --- |
| Cb | AV1 | R-G | 91 |
| Cb | LV1 | R-G | 91 |
| Cb | LV7 | S-P | 92 |
| Cb | RV7 | L-P | 87 |

**Supplementary table 2:** All mutations identified in the classic escape mutant selections by [Gerhard et al.
| antigenic site | mutant virus name | mutation | WSN HA numbering |
| --- | --- | --- | --- |
| Cb | LV7 | S-P | 92 |
| Cb | RV6 | E-K | 132 |
| Cb | RV7 | L-P | 87 |

**Supplementary fille 1:** This file shows the sequence and sequential A/WSN/1933 HA numbering scheme used in this study. It also gives the corresponding numbers in the 1RVX PBD HA structure and in an H3 numbering system.

**Supplementary file 2:** The computer code to perform the data analysis described in this manuscript is provided in the form of an iPython notebook.

**Supplementary file 3:** `ZIP` file showing logo plots spanning the entire gene illustrating the differential selection for each antibody at the concentration used in Figure 4. These data are the average across the replicate libraries for each antibody.

**Supplementary file 4:** `ZIP` file giving all mutation differential selection for each antibody at each concentration tested. These values are the average across the replicate libraries for each condition.

